# Regularization Improves the Robustness of Learned Sequence-to-Expression Models

**DOI:** 10.1101/393835

**Authors:** Bryan Lunt, Saurabh Sinha

## Abstract

Understanding of the gene regulatory activity of enhancers is a major problem in regulatory biology. The nascent field of sequence-to-expression modelling seeks to create quantitative models of gene expression based on regulatory DNA (cis) and cellular environmental (trans) contexts. All quantitative models are defined partially by numerical parameters, and it is common to fit these parameters to data provided by existing experimental results. However, the relative paucity of experimental data appropriate for this task, and lacunae in our knowledge of all components of the systems, results in problems often being under-specified, which in turn may lead to a situation where wildly different model parameterizations perform similarly well on training data. It may also lead to models being fit to the idiosyncrasies of the training data, without representing the more general process (overfitting).

In other contexts where parameter-fitting is performed, it is common to apply regularization to reduce overfitting. We systematically evaluated the efficacy of three types of regularization in improving the generalizability of trained sequence-to-expression models. The evaluation was performed in two types of cross-validation experiments: one training on D. melanogaster data and predicting on orthologous enhancers from related species, and the other cross-validating between four D. melanogaster neurogenic ectoderm enhancers, which are thought to be under control of the same transcription factors. We show that training with a combination of noise-injection, L1, and L2 regularization can drastically reduce overfitting and improve the generalizability of learned sequence-to-expression models. These results suggest that it may be possible to mitigate the tendency of sequence-to-expression models to overfit available data, thus improving predictive power and potentially resulting in models that provide better insight into underlying biological processes.

## Introduction

Enhancers [1], [2], also called cis-regulatory modules or ‘CRMs’ in some contexts, are ∼1 Kbp long sequences that harbor DNA binding sites for one or more TFs that act together to regulate a gene’s expression pattern [3]–[6]. Discovery of enhancer locations genome-wide and characterization of their regulatory activities are major problems in regulatory genomics today. A sequence-to-expression model (S2E model) is a function that maps an enhancer’s sequence to the regulated gene’s expression level in a cellular condition, given the relevant TF expression levels in that condition. It is thus an approach to the enhancer activity prediction problem. While current efforts at gene regulatory network (GRN) reconstruction [7]–[11] are dedicated primarily to identifying relevant regulatory inputs to a gene (and hence to its enhancers), an S2E model focuses on *quantitative* modeling, e.g., determining the input-output function at such a resolution that consequences of small changes to the inputs can be predicted, or explaining quantitative variations of a single gene’s expression across many cellular contexts. That is, S2E modeling builds upon the qualitative and discrete view afforded by GRNs, to provide quantitative predictions of gene expression.

One of the most promising paradigms of S2E modeling today is that represented by thermodynamics-based models [12]–[19]. The hallmark of these models is that they use the language of statistical thermodynamics to map molecular interactions involving proteins and DNA to gene expression levels. In previous work, authors have developed [19] and applied [20], [21] the thermodynamics-based model named ‘GEMSTAT’ to understanding the cis-regulatory code of developmental enhancers in Drosophila. GEMSTAT examines the three major components involved in regulating transcription: (a) DNA sequence (the enhancer), (b) TF molecules, and (c) the basal transcriptional machinery or “BTM”. It estimates binding site affinities from sequence using a position weight matrix (PWM) description of each TF’s binding specificity. It uses a single free parameter per TF to convert binding site affinities to their binding constants, and another free parameter to model the activation or repression strength of the TF. Thus, it uses only two free parameters per TF (and optional additional parameters for any cooperative binding mechanisms to be included in the model). This is in contrast to commonly used GRN reconstruction methods [22] that employ one free parameter per TF-gene pair and rely heavily on regularization to prevent over-fitting.

In recent work, Samee et al. [23] showed how GEMSTAT can be used to model expression data on a single gene (or enhancer), reveal underlying mechanisms at a quantitative level, and make accurate predictions about the effect of minor sequence changes such as mutating TF binding sites. Unfortunately, they found that even the modest number of free parameters in GEMSTAT (∼2 per TF, implying ∼10 parameters in a typical model using five TFs) leaves the data-fitting problem as largely unconstrained, opening the door for over-fitting. They addressed this problem by generating an *ensemble* of parameterizations (assignment of values to free parameters) that are consistent with the available data, rather than opting for the single best parameterization as is typically done [19] with such modeling approaches. Ensemble modeling of cis-regulatory sequences, as proposed in [23], embodies the view that the biologist investigating a gene’s expression control should be aware of *all possible explanations* of how the sequence encodes that control. Each parameterization of GEMSTAT is a possible explanation, which should be entertained until evidence to the contrary emerges from additional experiments [24].

In this work, we investigate a complementary approach to tackling the problem of over-parameterization in S2E modeling. We noted that the model ensembles reported by Samee et al. [23] often made erroneous predictions on distant orthologs of the enhancer sequences that they were trained on, a sign of potential over-fitting. We therefore explored different regularization techniques to soften the search topology of parameterizations when constructing an ensemble of S2E models from sparse available data. We adopted ‘noise-injection’ [25], L2 regularization, and L1 regularization, individually or in combination, and showed that the resulting ensembles of S2E models have greater predictive accuracy than those trained without regularization, when tested on unseen sequences.

## RESULTS

### Overview

The GEMSTAT model maps an enhancer’s sequence to target gene’s expression level in a set of cellular contexts, given the concentration levels of a fixed set of relevant TFs in those contexts. The model has free parameters that are fit to training data, which must include the inputs (sequence and TF concentrations) as well as outputs (target gene expression levels) of the model. The GEMSTAT implementation begins with an assignment of values to all free parameters, and optimizes it to improve the goodness-of-fit between model predictions and training data. We adopted the ensemble modeling approach of our previous work, where the numeric optimization of parameters is carried out multiple times, each time using a different initial parameterization. To systematically and quantitatively judge the utility of regularization for sequence-to-expression modeling, we implemented a workflow (Figure 1) as follows:

**Figure 1.**
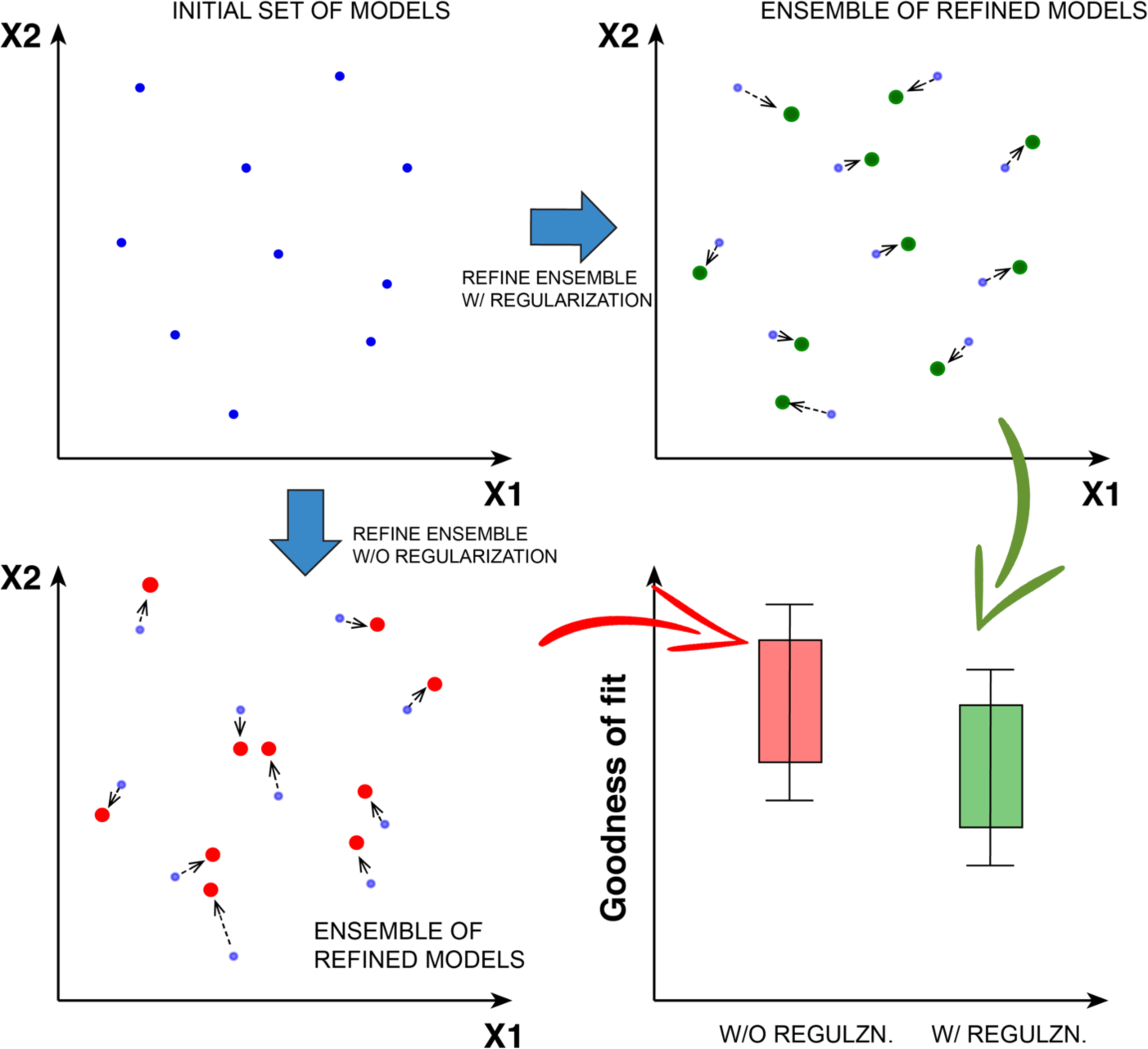
A schematic view of the process used to compare training with and without different forms of regularization. First, an initial set of model parameters is created randomly. Then, that set of parameters is used as starting points for model refinement under two different refinement methods. One is the traditional refinement method, and the other is the traditional method augmented with one or more forms of regularization. Finally, the goodness-of-fit values of the two ensembles of models are compared on held-out data to determine if either ensemble performs better with statistical significance.

1.A set of random initial parameterizations (points in the parameter space) are chosen, using appropriate ranges for each parameter, as selected in [23].

2.Each of these initial parameterizations is refined, i.e., optimized by GEMSTAT’s parameter fitting algorithm, both with and without regularization. Two ensembles of optimized parameterizations are thus obtained, differing only in the use of regularization during optimization. Parameter fitting is done using training data (enhancer sequence, TF concentrations, gene expression) from *D. melanogaster*.

3.Each parameterization in each ensemble is then used to predict the expression profile driven by a different enhancer, called the ‘test’ enhancer, with the goal of testing the model’s generalizability. TF concentration data used in making predictions are left unchanged in this test. A goodness-of-fit score is computed in the form of an ‘RMSE’ (root mean squared error) between the known expression profile driven by the test enhancer and that predicted by the model using that enhancer’s sequence.

4.The RMSE scores from these two ensembles of models are compared via a t-test.

In the remaining sections, we describe how we used the above strategy to demonstrate the advantages of using regularization during construction of GEMSTAT model ensembles. All of our tests involved enhancers that endogenously drive expression in a non-uniform ‘pattern’ along the dorso-ventral (D/V) axis of the early *Drosophila* embryo. The training and test data thus included TF concentration (input) and gene expression (output) levels at uniformly spaced points, called ‘bins’, along the D/V axis.

### Training with Noise Injection Improves the Cross-Species Predictive Accuracy of GEMSTAT Models

We reasoned that a robust GEMSTAT model ought to correctly predict the gene expression profile driven by an enhancer using the given TF concentrations profiles as well as slightly perturbed versions of the concentration profiles. This reflected our intuition that the ‘true’ model should not make drastically different predictions in the face of minor fluctuations in TF concentrations. Therefore, we modified the parameter fitting procedure by creating multiple copies of the training data set, injecting ‘noise’ into the inputs of all but one of these copies, and training models on the full collection of training data thus generated. We refer to this as training with ‘noise-injection’ [25], [26].

The data set modeled in this first test was the wild-type expression of the *ind* gene in the early *D. melanogaster* embryo. This developmental gene has a well-known enhancer that drives expression restricted to the neuroectodermal region along the D/V axis of the blastoderm stage embryo. The *ind* enhancer was the subject of extensive ensemble modeling in previous work [23], and is known to be regulated by the TFs Dorsal (DL), Zelda (ZLD), Twist (TWI), Snail (SNA) and Capicua (CIC), whose concentration/expression profiles along the D/V axis are also known (see Figure 2A and Methods).

**Figure 2.**
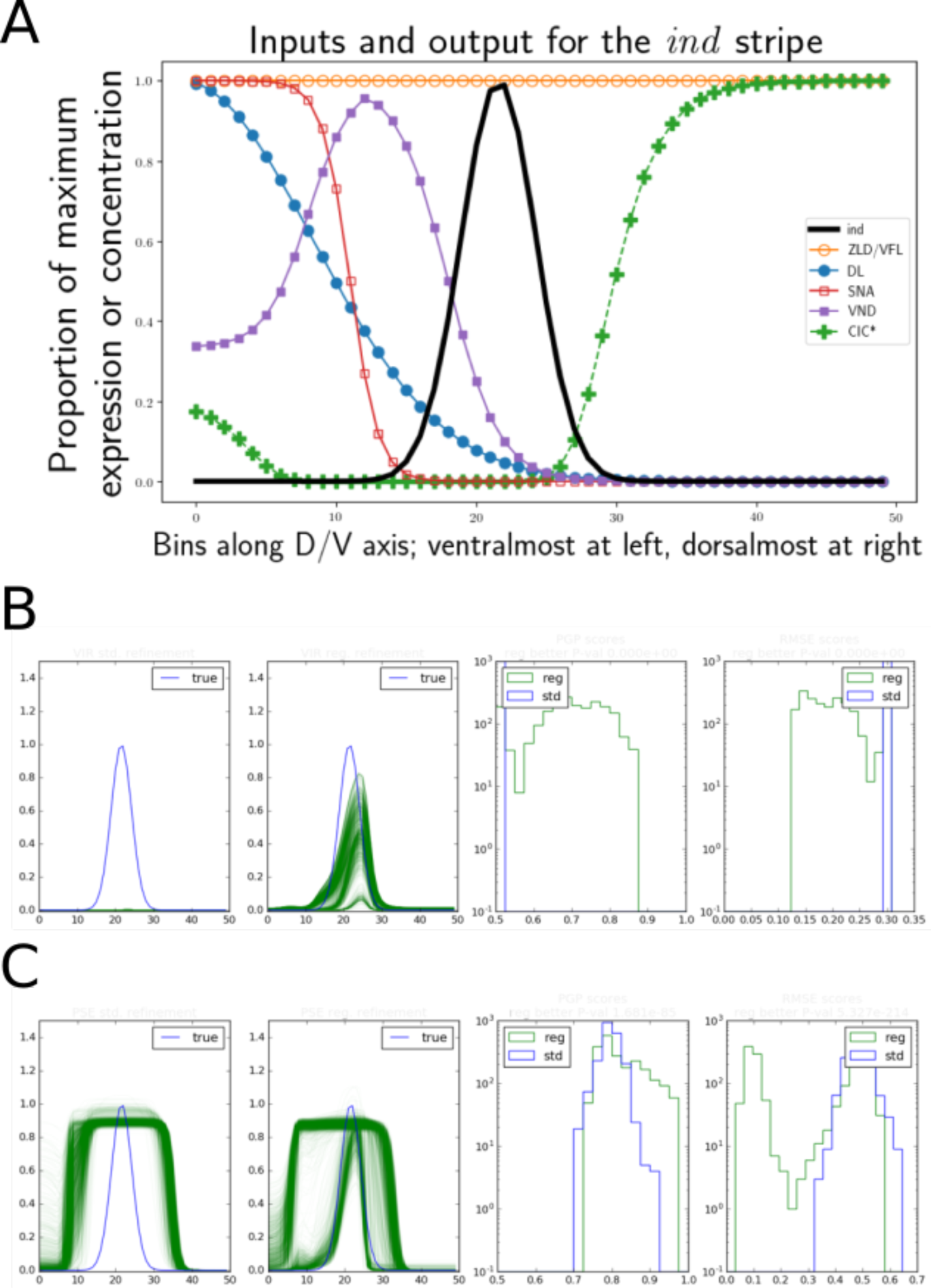
**(A)** All input and output curves from [23], used in evaluations involving the *ind* enhancer and reported in Tables 1 and 2. *Ind* expression displayed as the heavy solid line. Transcription factors are displayed as lines with markers. Effective CIC concentration (CIC*, plus signs) was calculated as described here and in [23] with a *cic_att* parameter of 16.0. **(B)** Example ensemble comparison for D.virilis, corresponding to Table 2, column 1. The first panel shows predictions (green) from models trained under standard refinement, with ground-truth in blue. The second panel shows predictions for models trained with noise-injection. The third panel displays a comparison of the histograms of PGP scores [20] for standard and noise-injected models. The fourth panel is the same comparing RMSE scores between the two ensembles. **(C)** Example ensemble comparison for D.pseudoobscura, corresponding to Table 2, column 2. Semantics are the same as panel B.

To evaluate the efficacy of noise injection for learning robust GEMSTAT models, we trained two ensembles of models – one with noise-injection and the other without – on the *ind* dataset from [23], following the workflow described in the previous section. Two thousand and one hundred initial parameterizations were randomly generated, and each was refined two ways using the GEMSTAT optimization procedure, either on the original training data, or on an expanded data set where the original is supplemented 20 noise-injected copies. The two resulting ensembles of optimized models were then compared for difference in their goodness-of-fit (RMSE) scores. This comparison was performed separately on the *ind* enhancer obtained from *D. melanogaster* as well as its orthologs from nine other *Drosophila* species. (Note that training data were exclusively from *D. melanogaster*, so evaluations on other species are on unseen data.) The first column in Table 1 gives the p-values from a Welch’s t-test used for these comparisons. As expected, the reduction of over-fitting resulted in worse fits on the training species, *D. melanogaster*, and the very closely related *D. simulans (not shown)*. On the more distant species, the ensemble of models trained with noise-injection significantly outperformed that of traditionally trained models for six of nine orthologs, was significantly worse for two orthologs, and statistically indistinguishable for one ortholog.

**Table 1:** Statistical evaluation of the effect of regularization during training of models. Ensembles trained on the *D. melanogaster ind* enhancer are evaluated on orthologs of the enhancer from each of nine other species (rows, sorted according to divergence times from *D. melanogaster*). ‘Better/worse’ indicates that an ensemble trained with a form of regularization has better/worse fits vs an ensemble of models trained without regularization. Shown in parentheses are p-values of Welch’s t-tests comparing RMSE (goodness of fit) scores of the two ensembles. Each of columns 1-3 evaluates a different form of regularization. **Column 1**: Noise regularization with N=20 copies and σ_0_ = 0.05was used to train 2100 models from random starting points as in [23], as described in the main text. **Column 2**: *L*_2_ regularization was used to train 2100 models. Analysis of noise-regularized models suggested that the scaling parameter contributed most to improvement, so we fixed it to 1.0 to avoid giving the regularized method a simple advantage in this regard. **Column 3**: *L*_1_regularization was tested in a test otherwise identical to that of column 2.

| | Noise-Injection | $L_2$ regularization | $L_1$ regularization |
| --- | --- | --- | --- |
| D. sechellia | Worse (0.324) | Better (4.6e-67) | Better (8.4e-39) |
| D. yakuba | Better (3.9e-88) | Better (1.6e-210) | Better (4.9e-210) |
| D. erecta | Better (3.1e-99) | Better (5.2e-86) | Better (5.1e-198) |
| D. ananassae | Worse (1.7e-48) | Better (6.0e-51) | Better (3.2e-97) |
| D. pseudoobscura | Better (3.3e-195) | Better (3.3e-87) | Better (1.2e-157) |
| D. persimilis | Better (1.8e-167) | Better (7.0e-93) | Better (7.8e-187) |
| D. mojavensis | Better (3.0e-04) | Better (8.7e-262) | Better (0.0) |
| D. virilis | Better (1.1e-70) | Better (4.2e-128) | Better (1.7e-222) |
| D. grimshawi | Worse (2.6e-10) | Worse (6.4e-118) | Worse (2.1e-264) |

**Table 2:** Comparison of ensemble obtained by noise-regularization versus ensembles reported in Samee et al. [23]. Evaluations and comparisons follow the same scheme as for Table 1 (also explained in text). ‘Better/Worse’ indicates that an ensemble trained with noise-regularization has better/worse fits, and p-values in parentheses are from Welch’s t-tests comparing RMSE scores for the two ensembles of models. **Column 1**: The final 2128 models from [23] serve as a baseline for evaluation of the 2100 models obtained from random starting points and refined by noise-regularization (same ensemble as that evaluated in Table 1 column 1). A p-value of 0 indicates that the p-value computed by the statistical software was smaller than its minimum possible p-value. **Column 2**: The final 2128 models from [23] were further refined for 1 epoch, with and without noise regularization, and the two resulting ensembles were compared.

|  | Ab Initio Noise-Injection vs. final ensemble from Samee et al. [23] | Refinement, with vs. without Noise-Injection, with ensemble from [23] as initialization |  |
| --- | --- | --- | --- |
| D. sechellia | Worse (0.0e+00) | Worse (0.0) |  |
| D. yakuba | Better (5.1e-109) | Better (4.0e-09) |  |
| D. erecta | Better (2.0e-209) | Worse (3.2e-09) | *1 |
| D. ananassae | Better (0.0) | Better (0.0) |  |
| D. pseudoobscura | Better (0.0) | Better (2.3e-234) |  |
| D. persimilis | Better (0.0) | Better (5.5e-233) |  |
| D. mojavensis | Better (0.0) | Better (5.8e-232) |  |
| D. virilis | Better (0.0) | Better (1.1e-234) | *2 |
| D. grimshawi | Better (insignificant) (0.947) | Worse (7.2e-150) | *3 |

We also sought to confirm that noise-injection during training generates more generalizable models compared to the ensemble of high accuracy models trained by Samee et al. [23]. The first column of Table 2 compares the 2100 models obtained by us using noise-injection (as above) against the 2128 best models reported by Samee et al. [23]. Performance was significantly better on nearly every ortholog except for the most closely related species, where it is expected to be worse (see Supplementary Figure S4). Results for *D. grimshawi* were not significantly different. In *D. virilis*, the ensemble of models from Samee et al. [23] predicted no expression at all, while most noise-trained models reproduce a correctly located stripe of *ind* expression (Figure 2B and Supplementary Figures S1 through S5).

In a related exercise, we took the ensemble of models from [23] and used them as initial parameterizations for one round of additional refinement, both with and without noise-injection. As shown in the second column of Table 2 performance was better with statistical significance for six of nine orthologs, which included five of the six most diverged species from *D. melanogaster*. This provides further evidence that noise-injection leads to models that are better able to predict the regulatory function of more distantly related test enhancers. Visually, the outputs of these models (Supplementary Figure S1) show that the models from [23], after refinement without regularization, tend to predict overly wide *ind* stripes. (This is also true of the models taken directly from that paper, without any regularization; Supplementary Figure S4.) For instance, see ensemble predictions in column 1 of Supplementary Figure S5, species *D. pseudoobscura* (‘PSE’) and *D. persimilis* (‘PER’). Predictions made by ensembles obtained with regularization also predict break into two classes, one of which fits the true expression pattern accurately while the other appears overly wide. On *D. grimshawi* it is very hard to see much difference in the two sets of predictions. This shows that noise-injection based regularization used alone can improve the generalizability of trained models.

### L1 and L2 Regularization also improve model generalizability

L1- and L2-regularization are two commonly used techniques, that help avoid over-fitting of models to small data sets. We evaluated these two regularization schemes in the same manner as noise-injection was evaluated above. That is, a set of randomly selected models was refined using that particular regularization scheme and goodness-of-fit scores were compared to those from refinement without regularization. The results are shown in Table 1, second and third columns. We observed that models fit without regularization often predict overly wide stripes or even ectopic expression for cross-validation species (Supplementary Figures S2, S3), while models refined from the same random starting points under regularization more often produce tighter stripes and less often predict ectopic expression. When using L2 regularization, 8 of 9 cross-validation tests showed a better distribution of RMSE scores with statistical significance. For L1, 7 of 9 tests showed significantly better performance for the ensemble trained with regularization. Intriguingly, models trained with either regularization scheme, as well as those trained using noise-injection, showed significantly worse prediction (compared to models from the default training procedure) on the *D. grimshawi* ortholog (Table 1, last row). This shows that L1 and L2 based regularization can be used to improve the generalizability of trained models.

### A combination of noise-injection, L1, and L2 regularization improves fitting for other dorsal/ventral patterning enhancers

We next tested the advantage of regularization during model-training using a different set of enhancers – those associated with four other D/V patterning genes present in the neurogenic ectoderm; *Rhomboid (rho)*, *Vein* (*vn)*, *Ventral Nervous System Defective* (*vnd)*, and *Brinker* (*brk*). Here, we trained models using one of these four enhancers and tested predictions on the other three, functionally related enhancers in the same species, rather than on orthologs of the training enhancer. At the blastoderm stage in *Drosophila* embryonic development, the four chosen enhancers are all regulated by the same set of patterning inputs, i.e., the TFs Dorsal (Dl), Twist (Twi), and Snail (Sna). Their patterns are mostly similar (Figure 3), with some offset, but their enhancer sequences are completely different. An important use of GEMSTAT, and sequence-to-expression modeling in general, is to generate models that not only predict accurately, but do so by gaining insight into the true biological process taking place. An ability to generalize to completely different sequences is more indicative of such a model than is the ability to make predictions on similar sequences, e.g., orthologs.

**Figure 3.**
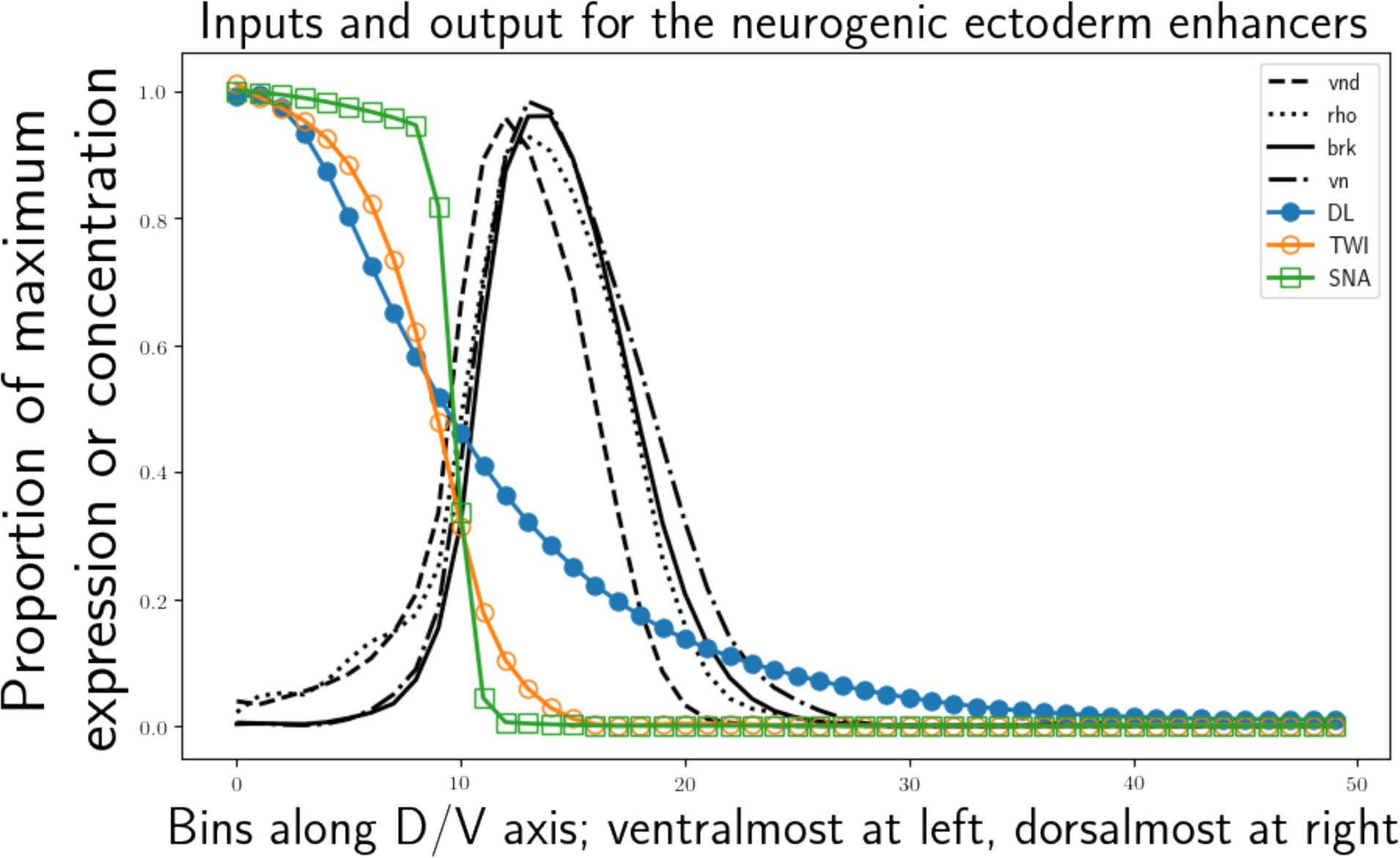
All input and output curves from [35], [36], processed as described in Methods, and used in evaluations reported in Table 3. Expression patterns are displayed as solid and dashed, markerless lines. Transcription factors are displayed with markers, according to the legend.

Table 3 shows the results of four separate training/cross-validation tests. In each, we trained GEMSTAT models on a single enhancer and compared the accuracy of predictions made by traditional versus regularization-trained ensembles on each of the other three enhancers. Noise-injection was used, with L1 regularization only used for cooperativity terms. This was because there is little prior knowledge of which TF pairs should be cooperative, and since L1 promotes sparsity, we should see extraneous cooperativities eliminated. Every test-case in Table 3 shows statistically significant improvement of results when using regularization. As can be seen in Supplementary Figures S6-S9, the improvements are often visually striking. In particular, predictions for the *rho* enhancer (with models trained on any of the other three enhancers) show a drastic improvement (see Supplementary Figures S7-S9, top row). At cross-validation time, traditionally trained models show a strong sensitivity to very small non-zero values of one input. This results in misplaced spikes in predicted *rho* expression, for nearly all of the traditionally trained models. These spikes are either strongly mitigated, or entirely absent in the predictions from regularization trained models. This shows that a combination of noise-injection, L1, and L2 based regularization can improve cross-validation to non-orthologous enhancers.

**Table 3:**
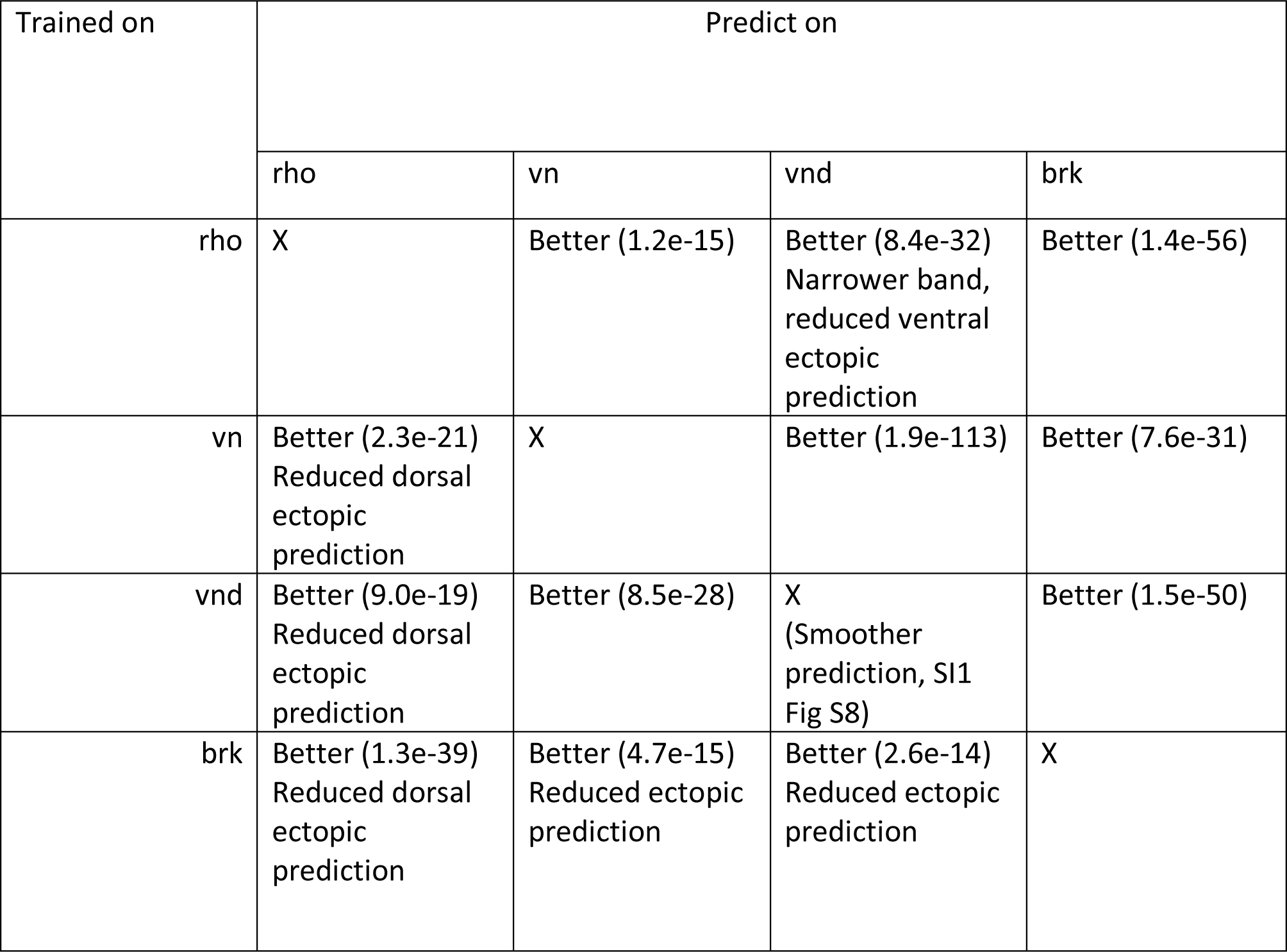
Comparison of ensembles obtained with and without noise-injection, using data on four D/V patterning enhancers in *D. melanogaster*. Shown in parentheses in each cell are p-values for Welch’s t-tests comparing RMSE scores of ensembles of 100 models trained on one enhancer and cross-validated on three other D/V enhancers. **Rows**: Two ensembles, one with and one without combined regularization, were trained on the enhancer listed in the ‘Trained on’ column. The RMSE scores of the two ensembles’ predictions on each cross-validation enhancer (‘Predict on’ columns) were compared with a Welch’s t-test, giving the p-values shown for the null hypothesis that the ensembles have identical performance. In all cases, the ensemble trained with regularization outperformed the traditionally trained ensemble with statistical significance.

## DISCUSSION

The goal of this research is to improve the way sequence-to-expression models are fit to data. That is a two-pronged task. First, we would like to improve the generalization accuracy of learned models. Learned models should be able to accurately predict the effects of mutation on sequences (cis-input), and the effects of unseen mixes of TF levels (trans-input). Second, we would like to improve the methods for model selection in the face of experimentally unknown interactions between players. This paper focuses mainly on the first point, though it begins to lay the groundwork for the second.

In Table 1 we present a basic evaluation of the two forms of regularization implemented here versus traditional model refinement. In Table 2 we evaluate our ensemble refinement method directly vis-`a-vis the final ensemble delivered by [23]. The comparisons reported in these two tables are based on model predictions on orthologs of the training enhancer. In contrast, Table 3 reports on comparisons based on cross-validation of enhancers within *D. melanogaster*, using hyperparameters decided upon in the previous tests.

All experiments resulted in marked improvements of generalizability. For the vast majority of cases, the ensemble refined with regularization outperforms the traditionally learned model with great significance. Tuning parameters (hyperparameters) for noise-injection proved to be relatively forgiving in the range of small values. Indeed, the first value we ever tried has turned out to be the best over several (not shown) experiments. Selection of L1 and L2 parameters was more difficult, and without enough data to perform a proper hyperparameter search, we settled on values small enough not to have drastic effects on the model, again in an intuitive way. The final set of experiments (Table 3) were run only once, with the hyperparameters decided upon in previous experiments. Not only did regularized models perform best in every case in this experiment, but in nearly every case a huge qualitative improvement is visually obvious.

With these three groups of experiments, we have shown strong evidence that improvement can be made in the way that sequence-to-expression models are fit to data. We took a fundamentally different approach to learning an ensemble of models than did Samee et al. [23]. In that work, the authors sampled millions of model parameter vectors, filtering for those that best fit the measured *D. Melanogaster ind* output. These were filtered, first for the 21000 models with the best RMSE scores on *D. Melanogaster ind*, and then to 2128 models that passed perturbation experiment filters. We suspect that the first filtering biases the models toward over-fitting the *ind* curve. Though Samee et al. reported that the models fell within 42 of their compartments in the model-parameter space, predicted curves for ortholog enhancers (SI, below) are all largely the same. This leads us to further suspect that the traditional model fitting problem is underspecified. Regularization offers a solution to the under-specification problem.

It may be noted that we discovered noise-injection ex-nihilo in an attempt to solve precisely the problems of ill-conditioned solution finding which the existing literature addresses. As a result, we have created a naïve implementation of noise-injection, itself an approximation to Tikhonov regularization. Further review of the literature reveals that the Levenberg-Marquardt non-linear least-squares optimization algorithm [27]–[31] directly implements Tikhonov regularization. In the future we hope to include this optimizer in GEMSTAT itself, doing away with noise-injection scripts.

## MATERIALS AND METHODS

### IND striping data

For *ind* (dorsal/ventral) [32] striping modeling, we took the datasets provided by the authors of [23], and used them without modification. This dataset includes curves for inputs *dl* (dorsal), *zld*/*vfl* (zelda/vielfaltig), *cic* (capicua), *sna* (snail), *vnd* (ventral nervous system defective); output *ind* (intermediate neuroblasts defective); and signaling kinase *dpERK* (doubly phosphorylated ERK [33], [34]) - all from late cell-cycle 14. Each curve had 50 bins along the ventral/dorsal axis, with bin 1 being ventralmost and bin 50 being dorsalmost. All curves were produced via experiments in *D. melanogaster*. The data is presented in Figure 2. The dataset also included *ind* enhancers from *D. melanogaster* and ten other Drosophilids. For orthologous enhancers, input and output patterns were presumed to match those of *D. melanogaster*.

### Neurogenic ectoderm striping system data

For other dorsal/ventral striping systems experiments, we collected data from http://dvex.org [35], [36]. While this website was not currently active when this research was performed, and has since been replaced with entirely different data, an archived versions of the website and original data are available from http://archive.org. The last useful snapshot being from 2009 (https://web.archive.org/web/20090408093453/http://www.dvex.org/). This dataset includes inputs *dl* (dorsal), *twi* (twist), and *sna* (snail); outputs *brk* (brinker), *rho* (rhomboid), *vn* (vein), and *vnd* (ventral nervous system defective), with outputs measured both for endogenous expression and expression of a minimal reporter driven only by the enhancer (not shown, available above). The database contains curves created by integrating the luminance over multiple stripes of confocal microscopy images, in addition to the individual bin values. Each image is registered to the *sna* gradient and endogenous *rho* mRNA [35], [36]. Each curve has 1000 points, from 0 at the ventral midline to 999 at the dorsal midline. In order to facilitate work at any number of D/V samples, we fit spline functions to those curves, with semi-manually selected distribution of knots, except for *dl* (discussed next). While every attempt was made to get splines that produced good curves, we did not force the curves to be perfectly smooth. This proved to be an important test of our method. Splines then allowed for the data to be up- or down-sampled to any desired number of bins.

In the case of *dl*, measured data does not cover the entire range of dl activity (there is a *dl* gradient from the ventral-most to dorsal-most points). Additionally, even for the coordinates where dl was measured, some of the tracks had missing data. To get an appropriate dl curve, we used a finite element differential equation solution that models production, diffusion, degradation, and the wraparound boundary implied by the 1-dimensional diffusion of *dl*. While technically it would be activating factors that are diffused through the perivitelline space [37], this approximation seems to fit the data well with only three parameters (effectively two, as at steady state, production and degradation must balance each other). The parameters of this diffusion model were fit with least squares to the region where data was available. The fit was nearly perfect, in contrast to the fit via a Gaussian curve used in [23] (not shown).

Enhancer sequences were taken from the RedFly database [38]. We used the enhancer “vnd NEE” for *vnd*, “rho NEE” for *rho*, “vn NEE-long” for *vn*, and “brk NEE-long” for *brk*. As reflected by their names, each of these sequences is known to drive expression during neurogenic ectoderm formation.

### Noise injection pre-processor

In order to realize noise injection without altering existing software, we implemented a tool that reads GEMSTAT input curves, copies the data bins, and applies noise. Output from this tool is in the standard GEMSTAT format, allowing unaltered versions of GEMSTAT to be used. Parameters to the tool are N, the number of copies to make of each bin (in addition to the original data); and σ_0_ and σ_1_, which control the noise. Each copied data point has Gaussian noise added, with standard deviation σ(y)=σ_0_ + σ_1_y, where y is the value of the curve in that bin. The noised input value is lower-bounded at 0.0. (Many early experiments, not shown, revealed that σ_1_ is unnecessary and may be set to 0.0.) For noise-injected training points, Gaussian noise with standard deviation 0.05 was added to normalized (max 1.0) input TF levels. Values falling below zero were thresholded to 0.0. Processed curves contain all of the original bins of the curve, augmented with N noised copies, thus for an input containing M bins, there will be (N +1)M output bins, NM of which have noise applied.

### Baking of effective *cic* levels

In [23], the authors calculated the effective concentrations of *cic* dynamically from the concentrations of dpERK, according to the following

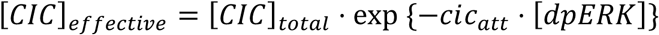

This results in small variations where *dpERK* levels are low causing very large variations in [CIC]_effective_. Our solution was to pre-calculate *cic*-attenuation before applying noise. We refer to this process as “baking” the *cic*-attenuation, or simply “baking”. Baked inputs can be handled by the base version (and the L1/L2 regularized version) of GEMSTAT, though it becomes impossible to optimize the *cic_att* parameter.

### Regularized GEMSTAT

We implemented L1 and L2 regularization in GEMSTAT. Some parameters can take separate regularization strengths, for example, scaling parameters (\beta) and cooperativities can be penalized separately from other parameters. The code is available at https://github.com/UIUCSinhaLab/GEMSTAT, currently in the ‘add_regularization’ branch, but will be merged to the ‘master’ branch in due time.

## ACKNOWLEDGMENTS

This work was supported by the NIH grant R01 GM114341 (to SS). We would like to thank Md. Abul Hassan Samee for providing his data and models. We also thank Farzaneh Khajouei for early technical assistance, enlightening discussions, and moral support.

